# Proteome-wide characterization and biomarker identification of intracranial aneurysm

**DOI:** 10.1101/725473

**Authors:** Tanavi Sharma, Keshava K. Datta, Munish Kumar, Gourav Dey, Aafaque Ahmad Khan, Kiran Kumar Mangalaparthi, Poonam Saharan, C. Shobia, Ashish Aggarwal, Navneet Singla, Sujata Ghosh, Amit Rawat, Sivashanmugam Dhandapani, Pravin Salunke, Rajesh Chhabra, Dalbir Singh, Aastha Takkar, Sunil K. Gupta, T. S. Keshava Prasad, Harsha Gowda, Akhilesh Pandey, Hemant Bhagat

## Abstract

The scientific basis of intracranial aneurysm (IA) formation, its rupture and further development of cerebral vasospasm remains incompletely understood. Deciphering the molecular mechanisms underlying these events will lead to identification of early detection biomarkers and in turn, improved treatment outcomes. Aberrant protein expression may drive structural alterations of vasculature found in IA. To unravel these aberrations, we performed untargeted, global, quantitative proteomic analysis of aneurysm tissue and serum from patients with IA. Samples were derived from patients of three clinical sub groups– 1) unruptured aneurysm 2) ruptured aneurysm without vasospasm 3) ruptured aneurysm who developed vasospasm. A total of 937 and 294 proteins were identified in aneurysm tissue and serum samples respectively. Several proteins that are known to maintain the structural integrity of vasculature were found to be dysregulated. ORM1, a glycoprotein, was significantly upregulated in both the aneurysm tissue and serum samples of unruptured IA patients. We employed a larger cohort of patients and validated ORM1 as a potential biomarker for screening of unruptured aneurysm using ELISA. Samples from ruptured aneurysm with vasospasm showed significant upregulation of MMP9 as compared to ruptured aneurysm without vasospasm. Using a cohort of ruptured aneurysm patients with and without vasospasm, we validated MMP9 as a potential biomarker for vasospasm. This study reveals pathophysiology underlying different clinical sub groups of IA and also suggests potential biomarkers.

## Introduction

Intracranial aneurysm (IA) is a pathological dilation of intracranial artery that is characterized by focal deterioration of the vessel wall that involves loss of internal elastic lamina, disruption of media and disintegration of adventitia and extracellular matrix of the artery (Frosen *et al.*, 2004, Nakajima *et al.*, 2000, Rajesh *et al.*, 2004). IA may be either familial or sporadic and is found to affect 3.2% of the general population (Vlak *et al.*, 2011). Patients who harbor IA are usually not detected and are manifested only once the aneurysm ruptures, leading to intracranial bleed and elevated intracranial pressure (Zoerle *et al.*, 2015). The consequences of rupture of an aneurysm are both on the cerebral and systemic physiology leading to multi system changes (Chen *et al.*, 2014). Despite technological advances in the management for prevention of rebleeding, the blood present in the intracranial space itself acts as a trigger for various inflammatory pathways leading to delayed cerebral ischemia and worsening the outcome of the patients or further adding to ongoing neurological insults (Archavlis and Carvi Y Nievas *et al.*, 2013, Chaichana *et al.*, 2010). Though the clinical spectrum of the disease has been established, mechanisms and pathophysiology of disease initiation and progression, i.e. aneurysm formation, rupture and subsequent development of cerebral vasospasm, are largely unknown until date.

There are no molecular biomarkers that can be used for early detection of intracranial aneurysm. Moreover, the patients at risk of aneurysmal rupture and those who can subsequently develop cerebral vasospasm cannot be predicted. A few targets for disease management have shown promise but have not been able to reduce the disease burden significantly. It is intriguing that despite various scientific and technological advances in research and clinical management, till date it has not been possible to reduce the burden of the disease which carries mortality of 30% and morbidity of 50% (Findlay *et al.*, 2016). Various studies directed towards deciphering the pathophysiology as well as to guide management of disease have not yielded beneficial results. Therefore, a more holistic understanding of the disease spectrum is the need of the hour. Patients who harbor IA are not victims of a single disease but a disease spectra that can range from an unruptured IA to aneurysmal subarachnoid hemorrhage (aSAH), hemorrhagic stroke and may further progress to cerebral ischemia (ischemic stroke). In this context, it is important to unravel the molecular basis of IA. Structural changes that are seen during aneurysm formation and rupture appear to be related to proteins as demonstrated in a few studies targeting single proteins (Connolly *et al*., 1997, Gaetani *et al*., 1997). Therefore, we envisaged a global, untargeted proteomic study of entire disease spectrum of patients with IA using both aneurysm tissue and serum samples. We employed a mass spectrometry (MS)-based quantitative proteomics approach to study protein based molecular alterations in patients with IA. To the best of our knowledge, this is the first report regarding whole proteome characterization of the entire disease spectrum in IA.

## Materials and Methods

The study was conducted at Post Graduate Institute of Medical Education and Research, Chandigarh and the Institute of Bioinformatics, Bangalore, India. The study was designed to unravel the proteomic differences in patients who harbor cerebral aneurysm in comparison to those who do not. Further, the study looked into the difference in protein expression between unruptured aneurysm and those who rupture and subsequently those who develop cerebral vasospasm. Consequently, four groups of patients were studied – a) control group (Group C), b) unruptured aneurysm (Group T1), c) ruptured aneurysm without vasospasm (Group T2) and d) ruptured aneurysm with vasospasm (Group T3). This study was approved by Institutional Ethics Committee PGIMER, Chandigarh (IEC-07/2015-268). Subjects were enrolled in the study only after obtaining an informed consent from the subject or their next of kin. Aneurysm tissue and blood samples were collected from the subjects of the four groups.

### Aneurysm-Tissue Samples

Aneurysmal wall tissue samples were collected from each test group (T1, T2 and T3) during surgery. For controls, (Group C), intracranial artery tissue was collected from the circle of Willis of those subjects undergoing autopsy and with no evidence of intracranial aneurysm. Samples were collected within 4 – 6 h following death. Tissue samples were collected in plain vials following washing with normal saline. Tissue samples were then stored at −80 °C until further use.

### Serum Samples

For test groups, blood samples were collected prior to surgery, before giving any anaesthetic agent, or any treatment. For controls, blood samples were collected from the subjects with negative angiography for any kind of cerebrovascular dysfunction. These were the subjects who underwent angiography for cerebrovascular evaluation and had no cerebrovascular disorder on angiography.

Blood samples (5ml) were collected in plain vials and were allowed to coagulate for 45 min at 37°C and centrifuged (3000 rpm for 15 min, 4°C) to obtain serum (supernatant). All serum samples were stored at −80 °C until further use.

### Study Design

Study was divided into two phases – Discovery Phase and Validation Phase.

#### Discovery Phase

In discovery phase, differentially regulated proteins were identified in aneurysm tissue and serum samples using untargeted high resolution mass spectrometry-based proteomics. We collected a total of 20 tissue samples and 20 serum samples for discovery phase analysis. Out of them, five tissue/serum samples were collected in each test group (T1, T2 and T3) and control group (C).

#### Validation Phase

In the validation phase, candidate biomarkers obtained from mass spectrometry data analysis were confirmed on a large cohort of samples. For validation phase, a total of 78 serum samples were collected and processed in four groups i.e. C, T1, T2 and T3. For screening marker of unruptured IA, 13 serum samples in Group C and 13 in Group T1 were collected. For validation of candidate molecule for vasospasm prediction, 26 serum samples in Group T2 and 26 in Group T3 were collected. For all the test groups, patients with unruptured IA (T1), ruptured IA without vasospasm (T2) and ruptured IA with vasospasm were diagnosed based on CT scan and angiographic examination. For controls, subjects with no evidence of intracranial aneurysm on autopsy or angiographic examination were included for the study.

### Sample preparation for LC-MS3 analysis

Mass spectrometry based quantitative proteomic analyses of aneurysm tissue and serum samples of groups C, T1, T2, and T3 were carried out as follows **(Fig. 1).**

**Figure 1:**
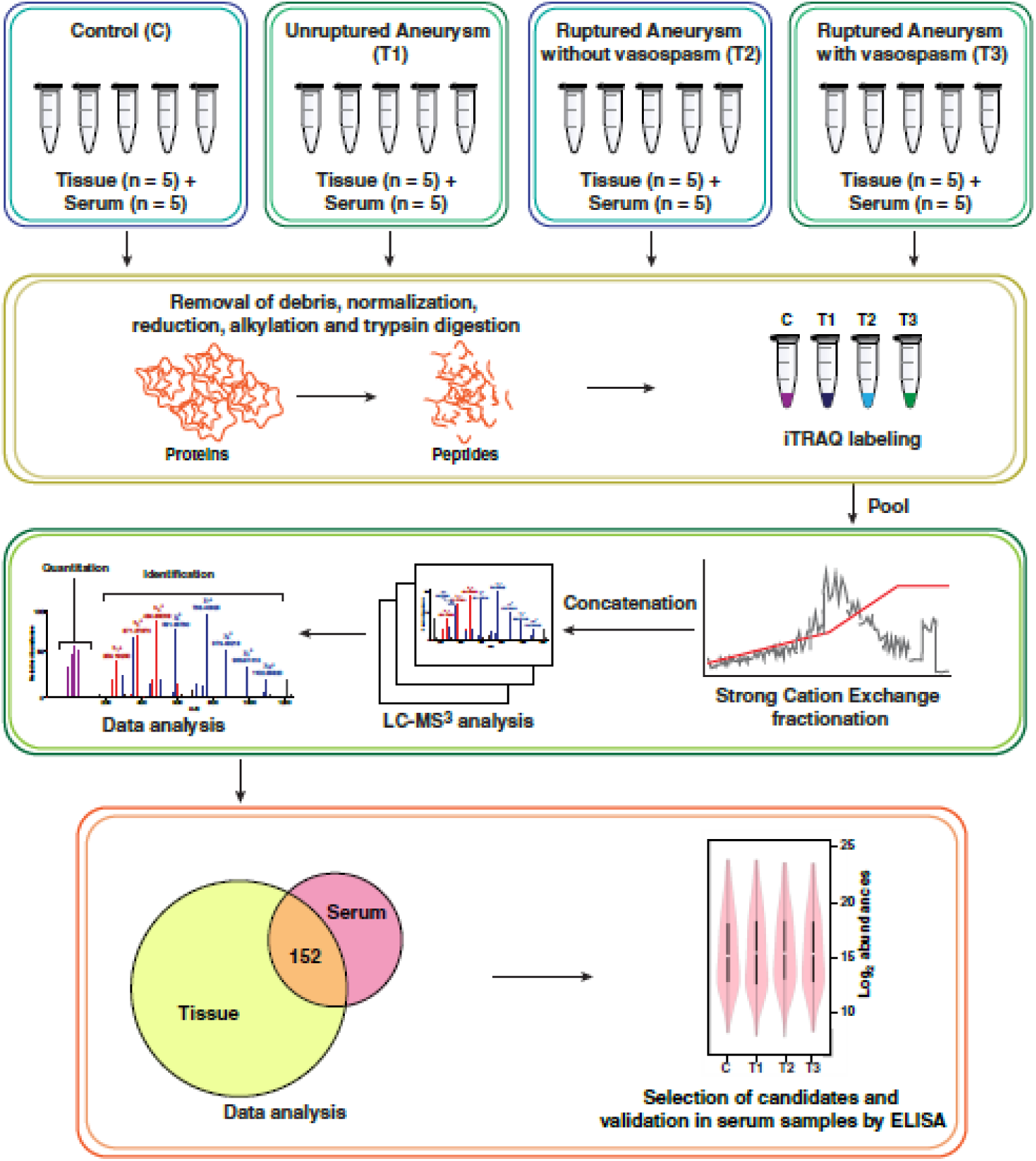
Overview of the study.

#### Protein Extraction

Equal masses of five tissue samples from each group i.e. C, T1, T2 and T3 were pooled and disrupted in liquid nitrogen using a hand-held disruptor. Subsequently, these were homogenized in lysis buffer (0.5% SDS, 100mM triethyl ammonium bicarbonate (TEAB)); centrifuged and clear supernatant was separated. Protein estimation was carried out using bicinchoninic acid (BCA) assay (Pierce, Waltham, MA;(Smith *et al*., 1985).

For serum samples, total protein was measured by BCA assay in groups C, T1, T2 and T3. Equal amount of proteins from five individual samples were pooled in each group. Serum samples were then depleted of highly abundant serum proteins using Human multiple affinity removal system–14 (MARS–14) column (Agilent Technologies, CA) as per manufacturers’ instructions.

#### Trypsin digestion

Equal amount of proteins from each group were reduced using Tris (2-carboxyethyl) phosphine (TCEP) at 60°C for 1h. Reduction was followed by alkylation using methyl-methanethiosulfonate (MMTS) at room temperature for 10min. Trypsin (modified sequencing grade; Promega, Madison, WI) was added (enzyme: protein = 1:20) to the samples and digestion was carried out at 37°C for 16h. Reaction was quenched by acidifying the tryptic peptides with formic acid.

#### iTRAQ labeling and fractionation

4-plex iTRAQ was used to label the peptides as per manufacturers’ instructions. Peptides obtained from Group C were labeled with isobaric group 114 while peptides for groups T1, T2, T3 were labeled with 115, 116 and 117 respectively **(Fig.1).** Labeled samples were pooled and subjected to strong cation exchange chromatography-based fractionation. A polystyrene-divinyl benzene copolymer modified with sulfonic acid Stage Tips (Empore Solid Phase Extraction Disk) were used as columns for fractionation. Sample digest containing iTRAQ labeled pooled peptides was reconstituted in 70 µl of 1% TFA and fractionated into 6 fractions with in – house prepared columns (Deeb *et al.*, 2015). The fractions were dried, reconstituted in 0.1% formic acid and desalted using C18 Stage Tips.

#### LC-MS3 analysis

iTRAQ labeled fractions were analyzed on Orbitrap Fusion Tribrid mass spectrometer (Thermo Scientific, Bremen, Germany) interfaced with Proxeon Easy-nLC II system (Thermo Scientific, Bremen, Germany). Each fraction was first loaded on a 2 cm long trap column packed with MagicC18AQ (Michrom Bioresources, Inc., Auburn, CA, USA). A linear gradient of 5% to 30% of solvent B (95% acetonitrile, 0.1% formic acid) at a flow rate of 300nl/min was used on an analytical column (75 μm x 20 cm, Magic C18AQ) for separation of peptides. Total run time was set to 120 min. Each fraction was analyzed three times. Mass spectrometer was operated on data dependent acquisition mode. Scan range was set on 400-1600 m/z. Orbitrap mass analyzer was operated at a mass resolution of 120,000 at MS level; For MS/MS and MS/MS/MS levels resolution was set to 30,000 and 60,000 respectively. Most intense precursor ions were selected and fragmented using higher energy collision dissociation (HCD).

#### Data Analysis

The data acquired were searched against Human RefSeq81 protein database. The searches were performed using SEQUEST and MASCOT through Proteome Discoverer (Version 2.1) software suite (Thermo Scientific, Bremen, Germany). The search parameters included trypsin as the proteolytic enzyme with one missed cleavage and oxidation of methionine as a dynamic modification. Static modifications were carbamidomethylation of cysteine and iTRAQ label at N-terminus of the peptide and lysine. Precursor and fragment ion mass tolerance were set at 10 ppm and 0.05 Da respectively. False discovery rate (FDR) was calculated using a target-decoy strategy and 1% FDR was applied at the peptide level. Quantification was carried out using the reporter ion quantifier node for iTRAQ available in Proteome Discoverer.

#### Data Availability

The mass spectrometry proteomics data have been deposited to the ProteomeXchange Consortium via the Pride partner repository with the dataset identifier PXD013442.

#### Enzyme linked immunosorbent assay (ELISA)

ELISA kits were procured from Cloud-Clone Corporation, USA. ELISA-based validation of selected candidate proteins was carried out as per manufacturers’ instructions. Briefly, 100µl of diluted serum samples, standard dilutions and blank were added to appropriate wells. ELISA plate was sealed with the help of sealer and incubated for 60 min at 37°C. Liquid was decanted off from each well after incubation. Detection reagent A (100 µl) was then added to each well. Again, the plate was sealed and incubated for 60 min at 37°C. Post incubation, wells were washed thrice with wash buffer and liquid was completely decanted off. 100µl of detection reagent B was added to each well and plate was incubated at 37°C for 30 min. The plate was washed three times with wash buffer. 90µl of substrate solution was then added and plate was incubated for 15 min at 37°C. 50µl of stop solution was added to each well for quenching the reaction. Absorbance was read at 450 nm using Infinite 200Pro plate reader, Tecan, Switzerland.

#### Statistical and Bioinformatics Analysis

For proteomics data, multiple t-tests with Holm – Sidak corrections were used for assessing the statistical significance between groups. Only those proteins that showed differential regulation with fold change cut-off of two and p≤0.05 were considered for further analysis.

For ELISA, unpaired t–test was used for assessing the statistical significance between groups (p≤0.05). Receiver Operating Curves (ROC) were also generated to assess sensitivity and specificity of the candidate proteins. All statistical analyses were carried out using GraphPad Prism version 6 (GraphPad Software, La Jolla California USA).

Principal component analysis (PCA) to check the variability in protein expression data between groups was performed using Array Track_HCA_PCA (Xu J *et al.*, 2010). Heat maps were generated by Morpheus software (https://software.broadinstitute.org/morpheus/) using hierarchical clustering method.

## Results

In a quest to decipher the pathophysiology of aneurysm formation and identify potential biomarkers for screening and differential diagnosis of intracranial aneurysm (IA), we performed iTRAQ–based quantitative proteomic analysis of IA wall tissue and serum samples of three groups of patients with a) unruptured aneurysm (T1) b) ruptured aneurysm (T2) and c) ruptured aneurysm with vasospasm (T3). Intracranial artery tissue samples and serum samples from subjects with no evidence of IA on autopsy or angiographic examination were used as controls (C).

### Demographic data

For discovery phase, we used 20 tissue and 20 serum samples in the four groups: C, T1, T2 and T3. For validating potential biomarkers, we included a large cohort of 78 patients (13 control subjects, 13 patients ofT1 group, and 26 patients each from T2 and T3 groups. Demographic details of the subjects recruited in the study are provided in **Table 1.**

**Table 1:** Demographic Data. (M= Male; F= Female, SD=Standard Deviation)

| Study Groups | Contol group (C) |  |  | Patients with unruptured aneurysm (T1) |  |  | Patients with ruptured aneurysm without vasospasm (T2) |  |  | Patients with ruptured aneurysm with vasospasm (T3) |  |  |
| --- | --- | --- | --- | --- | --- | --- | --- | --- | --- | --- | --- | --- |
| Phase | *Discovery Phase* |  | *Validation Phase* | *Discovery Phase* |  | *Validation Phase* | *Discovery Phase* |  | *Validation Phase* | *Discovery Phase* |  | *Validation Phase* |
| Type of samples | Tissue samples | Serum samples | Serum samples | Tissue samples | Serum samples | Serum samples | Tissue samples | Serum samples | Serum samples | Tissue samples | Serum samples | Serum samples |
| Sample size | n = 5 | n = 5 | n = 13 | n = 5 | n = 5 | n = 13 | n = 5 | n = 26 | n = 5 | n = 5 | n =5 | n = 26 |
| Age (Mean ± SD) | 31 ± 11 | 51 ± 8 | 43 ± 9 | 44± 16 | 49 ± 7 | 39 ±11 | 44 ± 15 | 49 ± 5 | 52 ± 13 | 43 ± 11 | 46 ± 8 | 50 ± 8 |
| Sex (M:F) | 5:0 | 4:1 | 8:5 | 2:3 | 2:3 | 7:6 | 2:3 | 2:3 | 11:15 | 2:3 | 2:3 | 11:15 |

*Values are mean ± SD or number of patients*

### Overview of proteomics data

In the past few years, quantitative proteomic analysis using isobaric mass tags has become the method of choice to compare proteomes of multiple physiological conditions. Here, we used iTRAQ, one such robust isobaric tagging method for quantitative proteomics of IA. To improve the depth and breadth of the study, we used both serum and tissue samples of IA. A total of 937 proteins were identified following tissue analysis. From serum samples, we identified 294 proteins. 152 proteins were commonly identified in tissue and serum samples. Detailed information of the identified proteins in tissue and serum samples is provided in **Supplementary Table 1 and 2** respectively.

Principal Component Analysis of both tissue and serum data **(Fig. 2)** showed a clear separation between the test groups and controls. While unruptured aneurysm (T1) samples clustered closer to the control group (C), ruptured aneurysms, both with and without vasospasm (T2 and T3), clustered on the same plane. However, we could observe complete separation between them.

**Figure 2:**
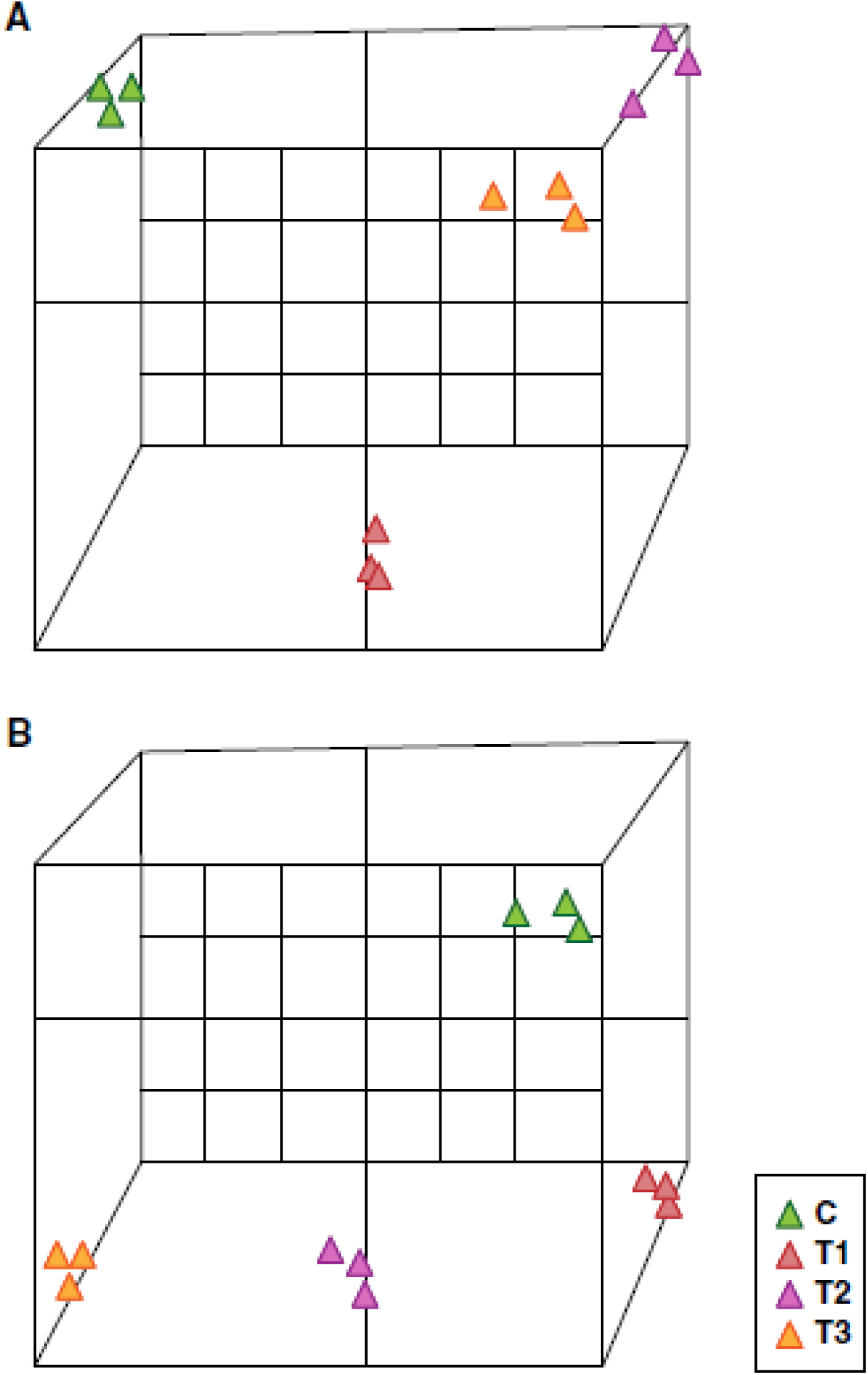
Principal Component Analysis. (A)Tissue Proteomics Data; (B) Serum Proteomics Data

We employed unsupervised clustering to identify molecular signatures associated with the clinical subsets of this study (**Fig. 3)**. We observed that the three clinical subsets were molecularly differentiated in both tissue (**Fig. 3A)** and serum (**Fig. 3B)** samples. Further, several molecules were found to be common in the signatures of tissue and serum. These molecules are described in the next sections.

**Figure 3:**
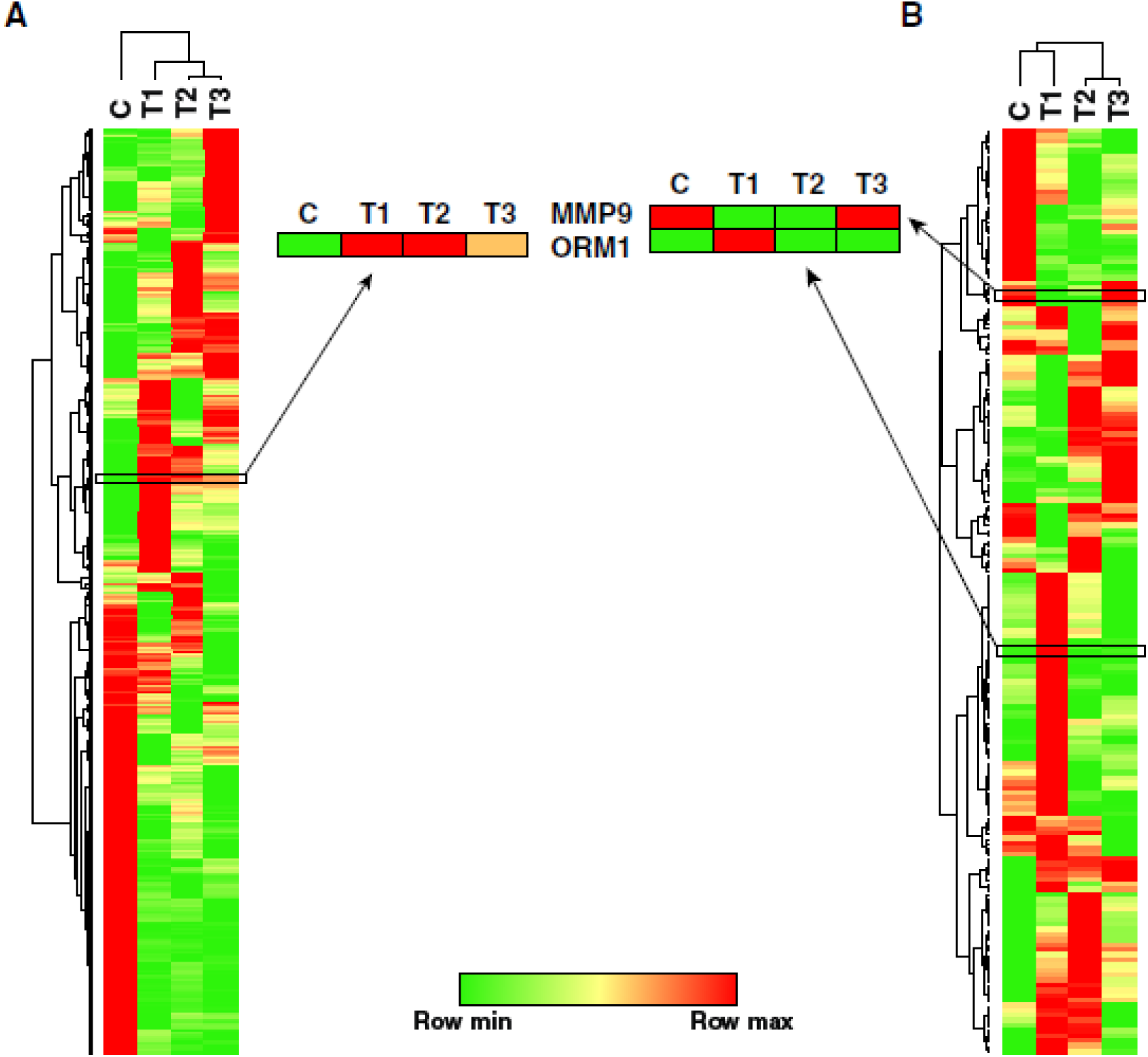
Heat map depicting molecular signatures associated with clinical subsets of patients with IA in **(A)** Tissue samples; **(B)** Serum samples

### Pathophysiology of aneurysm formation

To identify the molecular events that lead to the formation of IA, we studied the differentially regulated proteins in the IA tissue and serum samples of unruptured aneurysm-patients in comparison to healthy controls (T1/C). We identified 139 dysregulated proteins in tissue samples, 62 of which were upregulated and 77 were downregulated. In serum samples, 48 proteins were differentially expressed, out of which 30 proteins were overexpressed and 18 were downregulated.

Commonly dysregulated proteins between tissue and serum included α-1-acid glycoprotein 1 (ORM1), SAA2-SAA4 protein, α-1-antitrypsin (SERPINA1), serotransferrin (TF), apolipoprotein A-I (APOA1), transgelin-2 (TAGLN2), cytoplasmic actin (ACTG1), collagen alpha-1(XVIII) (COL18A1) and peroxiredoxin-1 (PRDX1).

Earlier studies have shown that inflammatory pathways and activation of the complement cascade are important factors during aneurysm formation (Taylor *et al.*, 2015, Turkmani *et al.*, 2015). Numerous proteins known to be involved in inflammatory processes and complement cascades were found to be overexpressed in both tissue and serum. Such proteins include α-2-macroglobulin (A2M) complement factor H (CFH), complement C3 (C3), complement C1q subcomponent (C1QC), complement C5 (C5), Azurocidin (AZU1), S100A4, and S100A9.

Thrombospondins are involved in maintaining the structure of vasculature by mediating cell-to-cell and cell-to-matrix interactions (Resovi *et al.*, 2014, Sherbet *et al.*, 2011). Thrombospondin1 (THBS1) has been shown to contribute to the development of Abdominal Aortic Aneurysm (AAA) by causing vascular inflammation through regulation of migration and adhesion of mononuclear cells (Liu *et al.*, 2015). Thrombospondin 4 (THBS4) is involved in local signaling in nervous system and also contributes to spinal sensitization and neuropathic pain state (Kim *et al.*, 2012). Role of THBS4 in inducing AAA has been recently demonstrated (Palao *et al.*, 2016). Our study, both THBS1 and THBS4 were overexpressed in tissue and serum respectively. Thus, thrombospondins may play a role in the formation of aneurysm.

### ORM1 is a potential screening biomarker for unruptured aneurysm

ORM1 is a glycosylated protein that has the capacity to bind and transport basic and neutral molecules(Fournier *et al.*, 2000). ORM1 is primarily synthesized by the liver but can also be produced in extrahepatic sites (Luo *et al.*, 2015). ORM1 is an acute phase inflammatory protein which also plays a role in injury induced angiogenesis (Ligresti *et al.*, 2012). In the present study, ORM1 was found to be 22 fold overexpressed in IA tissue and 9 fold upregulated in serum and thus was selected as a candidate molecule for validation.

ELISA-based validation of ORM1 was carried out in serum samples of patients with unruptured aneurysms (n = 13) and controls (n =13). We observed that levels of ORM1 were significantly higher (p= 0.004) in serum of unruptured aneurysm as compared to controls. The area under the curve (AUC) was found to be 0.78 with sensitivity and specificity of 76.92% and 81.92%, respectively (Fig. 4). Thus, ORM1 may be a potential biomarker for population screening of IA.

**Figure 4:**
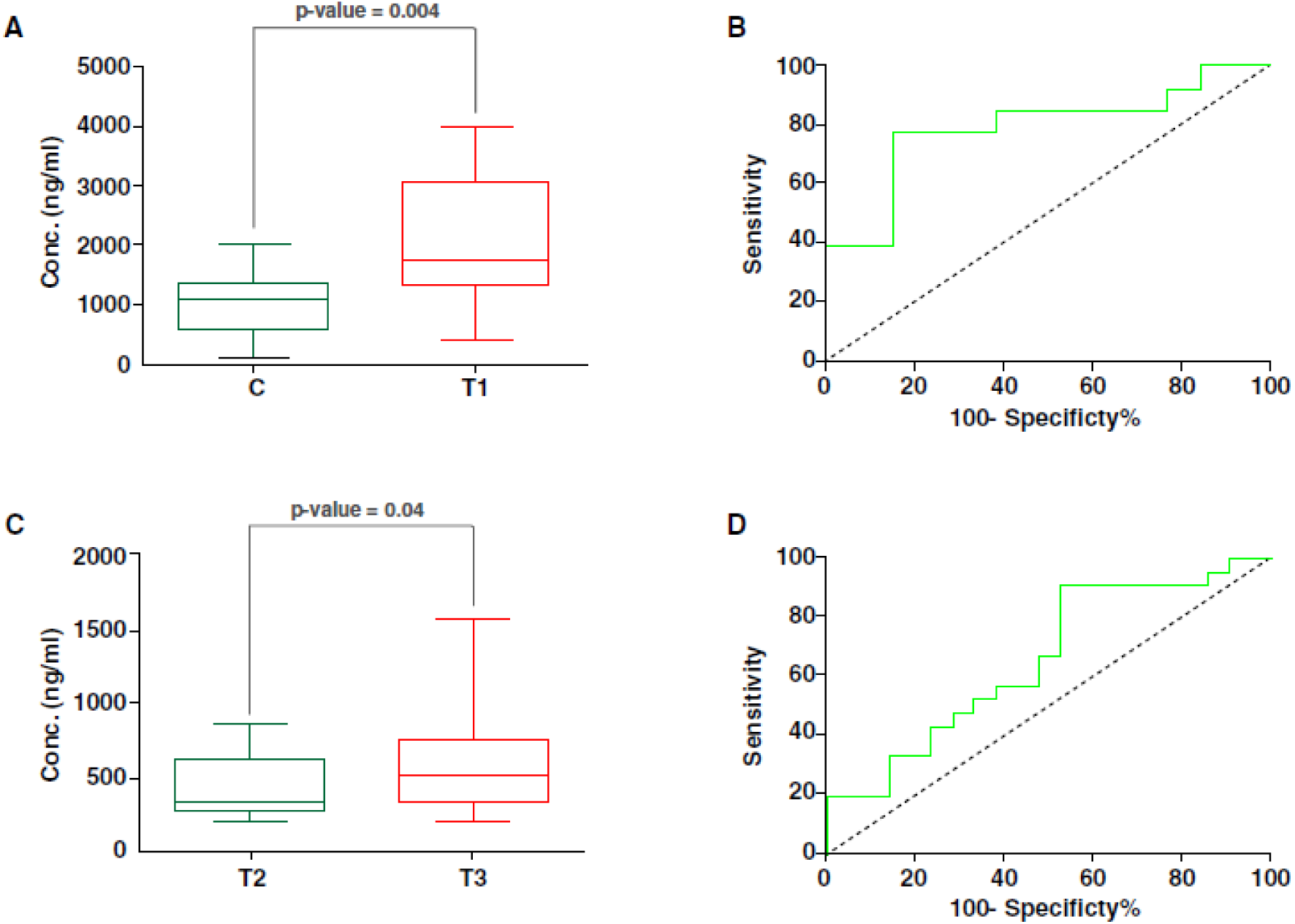
Estimation of ORM1 and MMP9 expression by ELISA. **(A)** Box plots depicting the difference in the expression level of ORM1 in patients with unruptured aneurysms compared to controls and **(C)** MMP9 in patients with ruptured aneurysms without vasospasm compared to ruptured aneurysm with vasospasm. (**B)** Receiver operating curve showing the specificity and sensitivity for ORM1 and **(D)** MMP9

### Molecular mechanisms underlying aneurysmal rupture

A significantly high number of patients with IA suffer from aneurysmal rupture every year. Comparing protein expression profiles of the samples with ruptured aneurysms with unruptured aneurysms is likely to provide an insight into molecular mechanisms underlying IA rupture. We identified42 differentially regulated proteins in tissues amples, out of which 28 were found to be upregulated and 14 were found to be downregulated. We could identify 37 differentially expressed proteins in serum samples, 10 proteins were overexpressed while 27 were found to be downregulated.

Chronic inflammation in aneurysm rupture has been associated in earlier studies (Hashimoto *et al.*, 2006, Hudson *et al.*, 2013). We observed overexpression of proteins related to inflammatory pathways in both tissue and serum samples. CRP was found to be upregulated in our study which has been demonstrated to be associated with an increase in size of AAA and its rupture (Turner *et al.*, 2015, Vainas *et al.*, 2003). Similarly, there was overexpression of SAA1 protein which could lead to aneurysmal rupture (Webb *et al.*, 2015).

### Proteomic aberrations leading to vasospasm

Development of vasospasm is seen to be detrimental to favorable prognosis post-aneurysmal rupture (Findlay *et al.*, 2016). Proteins that play a role in the formation of vasospasm may be employed as potential prognostic biomarkers. To identify such molecules, we compared protein expression in tissue and serum samples of patients with ruptured aneurysm with vasospasm and ruptured aneurysm without vasospasm (T3/T2).We identified 33 differentially regulated proteins in tissue samples, out of which 13 were found to be upregulated, while 20 were downregulated. While in serum samples, 36 proteins were found to be differentially expressed out of which 20 were overexpressed and 16 were showing downregulated expression.

To the best of our knowledge, this is the first proteomic study that compares differential expression of proteins in ruptured aneurysm with vasospasm and ruptured aneurysm without vasospasm. Superoxide has been reported to be associated with vasospasm (Suzuki *et al.* 1994; Mori, Nagata *et al.* 2001).We observed overexpression of protein Cytochrome b-245 light chain (CYBA) in tissue data. CYBA is known to associate with NOX3 to form a functional NADPH oxidase, which leads to generation of superoxide (Ueno, Takeya *et al.* 2005).

We also observed overexpression of proteins related to oxidative stress in patients with cerebral vasospasm. We found Myeloperoxidase (MPO) to be upregulated in tissue samples which has been previously found to be associated with acute ischemic stroke (Cojocaru *et al.*, 2010). Monoamine oxidase A (MAOA) was also found to be overexpressed. This protein has been demonstrated to oxidize biogenic amines such as norepinephrine and 5-hydroxytryptamine causing oxidative stress (Youdim *et al.*, 1984). In serum data, we observed elevated expression of peroxiredoxin-2 (PRDX2). This protein is related to oxidative stress and was found markedly elevated in the brain extracellular fluid in patients with acute ischemic stroke (Dayon *et al.*, 2011).

### MMP9 is a potential biomarker for vasospasm

MMPs are a family of extracellular and membrane-bound proteases capable of degrading or proteolytically modifying the ECM through interactions with collagenases, laminins and proteoglycans (Löffek *et al.*, 2011). MMP9 is a type IV collagenase that is involved in the cleavage of a variety of substrates on the cell membrane and extracellular matrix. MMP9 was found to be 13.4 fold upregulated in serum samples of patients with ruptured aneurysm with vasospasm as compared to that without vasospasm and was selected as a candidate molecule for validation.

ELISA-based validation of MMP9 was carried out using serum samples from patients with ruptured aneurysm with vasospasm and without vasospasm (n=26 in each case). Levels of MMP9 were significantly higher (fold change 1.4, p<0.05) in case of patients with ruptured aneurysm with vasospasm as compared to patients with ruptured aneurysm without vasospasm. The area under the curve (AUC) was found to be 0.77 with sensitivity and specificity of 66.67% and 71.43%, respectively **(Fig. 4).**

## Discussion

Over the past decade, MS based quantitative proteomics has emerged as the method of choice to identify molecular signatures of disease and to optimize their management. In succession to the same, a number of studies following MS based analysis from literature have provided new subset of protein biomarkers with possible therapeutic targets for IA (Jiang *et al.*, 2018, Wang *et al.*, 2015, Wang *et al.*, 2016, Xu *et al.*, 2015). Presently, there is no molecular study to our knowledge, depicting the differential expression of the proteins between three subgroups of patients with IA. Thus, to decode possible causative proteins and to understand the pathophysiology of aneurysm formation, rupture and the progression to vasospasm, mass spectrometry based analysis was performed. We analyzed aneurysm-tissue and serum samples of patients with IA using MS based quantitative proteomics. We chose to analyze both tissue and serum samples to look for the reflection of dysregulated tissue proteins (microenvironment of disease) in the serum samples (macroenvironment of disease). This would help us in identifying biomarkers in serum samples which is an easily accessible bio-fluid. This study has resulted in the identification of numerous clinically interesting proteins which may have significant contribution in the genesis and progression of the disease.

Despite being a dreadful complication of brain vasculature, the pathogenesis of IA formation remains poorly understood. Although the presence of IA is significant in general population (Vlak *et al.*, 2011), we are still at loss in detecting such patients prior to aneurysm rupture. Thus, it is important to study the biology of aneurysm formation as it may decipher molecules that can serve as biomarkers for population screening.

We studied differential expression of proteins in tissue and serum samples of patients with unruptured aneurysm and compared it to those of control samples. There were molecules such as ORM1, SERPINA1 and SAA2 – SAA4, which were commonly overexpressed in both tissue and serum data. Largely these have a role in the inflammatory pathway and inflammation has been demonstrated as an important pathological factor for formation of IA (Chalouhi *et al.*, 2012, Hosaka and Hoh *et al.*, 2014).

In our study we found upregulation of Matrix metalloproteinase (MMP-2) in serum data. MMP-2 is known to cleave the components of the ECM. Upregulation of MMP2 has been reported earlier (Maradni *et al.*, 2013). Further, Intercellular adhesion molecule 1 (ICAM1) is expressed on the endothelial cells. This protein binds to integrins and is involved in cell adhesion. ICAM1 was shown to promote experimental aortic aneurysms through recruitment of circulating leukocytes (Xu *et al*., 2016). In the present study (ICAM1) was also found to be 3.8 fold upregulated in serum data. Thus our findings are in concordance with the previous reports (Davis *et al.*, 1993, Szekanecz *et al.*, 1994).

Proteins that are crucial for organizing cytoskeleton and its maintenance were found to be down regulated e.g. myosin-11 (MYH11), filamin-A, (FLNA), actin, aortic smooth muscle (ACTA2), smoothelin (SMTN), α-actinin-4 (ACTN4), Tropomyosin1 (TPM1), tubulin α-1C chain (TUBA1C), laminin subunit beta-2 (LAMB2) and α-actinin-1 (ACTN1). They play a significant role in cell to cell communication, interaction and regulation of movement. Their altered expression may contribute to IA formation by diminishing the structural architecture of cerebral vasculature. Proteins related to actin cytoskeleton may result in reduced tensile stretch exerted by blood pressure, leading to formation of aneurysm. Filamin-A is an actin binding protein involved in cytoskeleton remodeling. It promotes mitosis by binding to cdc25C (Telles *et al.*, 2011). It is known to promote cell motility and migration. Down regulated expression of this protein demonstrates impaired division and growth of cells. Downregulated expression of proteins laminin subunit beta-2 (LAMB2) and α-actinin-1 (ACTN1) was observed in a recent study on IA (Wang *et al.*, 2016). Further, we found proteins such as Nidogen 1 (NID1), Cadherin 2 (CDH2) and Multimerin 2 (MMRN-2) to be downregulated in serum samples. These proteins play important roles in cell adhesion and growth (Galvagni *et al.*, 2017, Maitre and Heisenberg *et al.*, 2013, Miosge *et al.*, 2001). NID1 is a basement membrane glycoprotein which interacts with other components of basement membranes and plays a role in cell interactions with the extracellular matrix. Multimerin 2 (MMRN2) is an ECM glycoprotein that is involved in angiogenesis and growth. Cadherins protect the cells from mechanical shear and stress and its deficiency may result in the loss of the protective ability against wall shear stress (WSS). WSS has been reported to play a role in the formation of IA (Chatziprodromou *et al.*, 2007). Down regulation of the proteins related to cytoskeleton have also been reported in AAA formation (Modrego *et al*., 2012). This could explain the thinning of muscular layer of the intracranial vessel wall at the site of aneurysm formation.

Quantitative proteomic profiling of IA tissue and serum samples in patients with unruptured aneurysm when compared to controls enabled the identification of ORM1 as a potential biomarker for aneurysm formation. ELISA – based validation of ORM1 also confirmed its upregulation in serum of patients with unruptured aneurysm as compared to controls. ORM1 plays an important role in injury induced angiogenesis (Ligresti *et al.*, 2012). At the site of vascular injury, endothelial cells on the luminal side are persuaded by TNF-α (secreted from macrophages) to secrete ORM1. ORM1 negatively inhibits the TNF-α at the site of injury acting as an anti-inflammatory protein. At the same time it induces angiogenesis. A well-known pathophysiological consequence of injury induced angiogenesis is that it increases permeability of intercellular endothelial cell junctions leading to edema and extensive injury to the surrounding tissue (Weis and Cheresh *et al.*, 2005). Thus, ORM1 may play a role in aneurysm formation by inducing angiogenesis at the site of intimal injury that may progressively damage the vessel wall tissue **(Fig. 5).**

**Figure 5:**
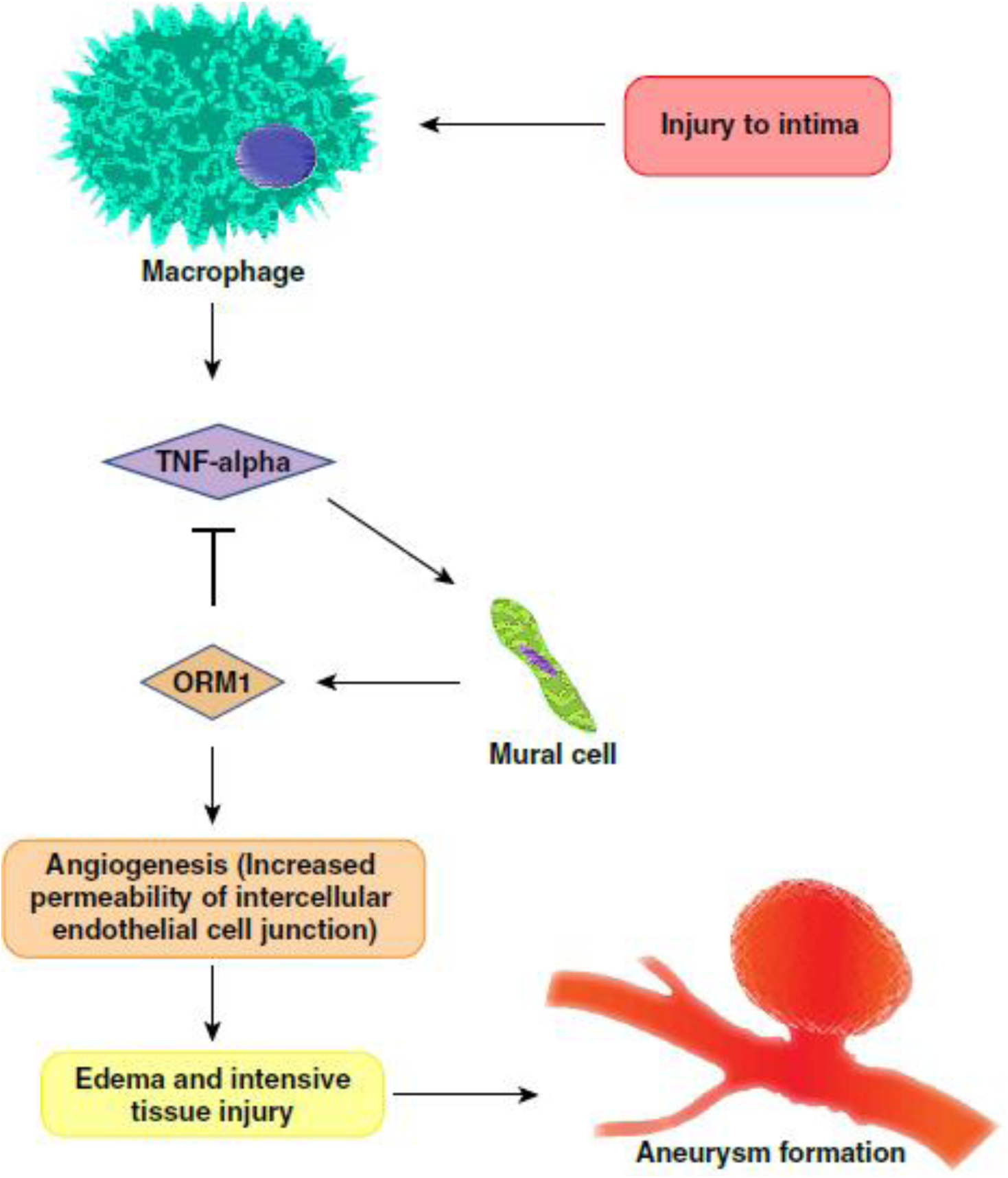
Potential mechanism of aneurysm formation through ORM1.

There has been lot of enthusiasm in trying to understand how aneurysms formed in the cerebral vasculature eventually progress to rupture. It is intriguing why only few of the aneurysm rupture and most of them do not. To understand the biology of aneurysmal rupture, we studied differential expression of proteins in tissue and serum samples of patients with ruptured aneurysm and compared it to those with unruptured aneurysm. We observed over expression of Cathepsin B (CTSB, a lysosomal cysteine protease) in the tissue samples. It is possible that an increased activity of CTSB in the tissue samples may contribute to aneurysm progression through caspase induced apoptosis of Vascular Smooth Muscle Cells (VSMCs), as supported by previous reports (Guo *et al.*, 2016, Aoki *et al.*, 2008). Interestingly, we also observed upregulation of Cystatin B (CSTB, a cysteine protease inhibitor) in the tissue samples. A positive association of cystatins and abdominal aneurysm size has been shown earlier (Wang *et al.*, 2018). Our observation regarding the overexpression of fibrinogens (FGA, FGB and FGG) in the tissue samples is in good agreement with the report of Al-Barjas*et al.* (2006), in which an increased level of fibrinogens was shown as a marker for aneurysm progression in AAA (Al-Barjas *et al.*, 2006). Various proteins involved in maintaining vascular integrity and haemostatic balance in the cell including Microfibrillar-associated protein 4 (MFAP4), Decorin (DCN), Tropomyosin family proteins (TPM1, TPM2 and TPM4) along with F-actin-capping protein (CAPZA1) were found to be upregulated in the tissue samples, deregulation of which was found to be associated with aneurysm formation (Modrego *et al.*, 2012).

On comparing the protein expression profiles of patients with ruptured aneurysm with and without vasospasm (T3/T2), we observed overexpression of AZU1 in the tissue samples. AZU1, an inflammatory cascade protein, is a component of neutrophil azurophilic granules and is implicated in proteolysis, apoptosis and phagocytic neutrophil oxygen independent killing (Soehnlein and Lindbom *et al.*, 2009, Watorek *et al.*, 2003). AZU1 is expressed in endothelial cells favoring contraction and increased endothelial permeability (Gautam *et al.*, 2001). Narrowing of blood vessels which lead to cerebral vasospasm is characterized by prolonged abnormal contraction of VSMCs (Chaichana *et al.*, 2010, Findlay *et al.*, 1991, Ladner *et al.*, 2013). Thus, an increased expression of AZU1 points to the fact that contraction of blood vessel may lead to cerebral vasospasm. S100A8 and S100A9 are the calcium binding proteins, which are known to play an important role in the regulation of inflammatory processes and immune response. Overexpression of these proteins also indicates the importance of inflammation in the development of vasospasm. Proteins related to cell adhesion and remodeling of the actin cytoskeleton such as integrin family proteins (ITGB3, ITGA2B and ITGB3), neutrophil defensin 1 (DEFA1), myeloblastin (PRTN3) and LTF (lactotransferrin) were found to be upregulated in patients with vasospasm. Blocking of integrins by their receptor antagonist GRGDSP has been shown to prevent vasospasm in earlier studies (Pradilla *et al.*, 2004),which is correlated well with our finding.

Lysosomal related glycoprotein (LAMP1) and Transforming growth factor-beta-induced protein (TGFBI) were found to be overexpressed in serum samples of patients with cerebral vasospasm in the present study. LAMP1and cathepsins activate the caspase cascades that induce apoptosis in the endothelia lining the intima (Haka *et al.*, 2016, Li *et al.*, 1998). Apoptosis in the endothelium results in its detachment which exposes the collagen of the internal elastic lamina, which promotes platelet adherence and thrombus formation (Smith *et al.*, 1985). Damage to endothelial cells also leads to reduction in nitric oxide production which demolishes the balance of vascular tone(Hino *et al.*, 1996). This exposed arterial state results in direct contact of SMCs with vasoactive agents, such as ATP in the blood stream, which causes constriction of vessel wall (You *et al.*, 1997). All these events may result in further narrowing of lumen resulting in cerebral vasospasm. TGFBI acts as a ligand recognition sequence for many integrins. This protein plays a role in cell-collagen interactions. Integrins play a role in altering calcium dynamics and phenotypic modulation of VSMCs (Balasubramanian *et al.*, 2007, Qin *et al.*, 2001). Thus, dysregulated expression of TGFBI may result in altered calcium dynamics leading to vasospasm.

Proteins involved in degradation of ECM such as matrix metalloproteinase (MMP9) were found to be significantly upregulated in serum data. One of the distinguishing features of cerebral vasospasm is prolonged contraction of the SMCs. Actin cytoskeleton of SMCs has a dynamic structure that plays an integral role in regulating the development of mechanical tension leading to contraction/dilation of smooth muscle tissues. Messed up actin polymerization has been linked with vasospasm in human saphenous vein (Hocking *et al.*, 2016). In the present study, downregulation of proteins related to cytoskeleton remodeling such as ACTA2, TPM2 and MFAP4, demonstrates their role in the development of vasospasm. Neuroserpin (SERPINI1), an inhibitor of tissue plasminogen activator (tPA) provides protection against cerebral vasospasm by preventing N-methyl-D-aspartic acid-induced neurotoxicity, which results in reducing tPA-mediated inflammation and disruption of the blood-brain barrier (BBB) (Gelderblom *et al.*, 2013, Lebeurrier *et al.*, 2005, Rodriguez *et al.*, 2008). Thus, downregulated expression of this protein in our study demonstrates its significance in pathology of cerebral vasospasm.

Quantitative proteomic profiling enabled the identification of MMP9 as a potential prognostic biomarker for vasospasm formation and was validated on a larger cohort of patients using ELISA. MMP9, a matrix metalloproteinase has been implicated in the pathophysiology of blood brain barrier (BBB) disruption and cerebral edema (Obermeier*et al.*, 2013, Turner and Sharp *et al.*, 2016). The mechanism is complex, but on disruption of BBB, there is leakage of the inflammatory mediators which leads to arterial dysregulation due to leukocyte migration. This blockage in the artery may lead to its narrowing, resulting in vasospasm. Further, in a mouse model, protective role of minocycline, an inhibitor of MMP9, for cerebral vasospasm was demonstrated recently (Vellimana *et al.*, 2017). Overall, it can be concluded that MMP9 has the potential to be used as a prognostic biomarker as well as a therapeutic target.

There are some limitations to the current study. We had a small sample size of patients in discovery phase. This was due to the difficulty in obtaining tissue samples of aneurysm wall. However, validation was done in a larger cohort based on sample size calculation. Nonetheless, it would be prudent to carry out further studies using a larger patient population. Although we have identified ORM1 and MMP9 as potential biomarkers, they need to be tested in a larger, geographically distinct cohort of patients to determine their diagnostic potential.

In spite of these variations, this study provides the first whole proteome characterization of the entire spectra of the disease in IA-tissue - and serum samples. This study indicates multiple proteins that may have a role in the pathogenesis of aneurysm formation, rupture and subsequent development of cerebral vasospasm. Findings from this study could lead to better diagnostic capabilities and therapeutic interventions.

## Acknowledgements

The authors thank the patients and family members for their participation in the study. We would like to express our greatest gratitude to Late Dr. K.K. Mukherjee and Late Dr. V. K. Grover for their invaluable guidance and support for the conduct of the study.

## Competing interests

The authors declare that they have no competing interests.

## Funding

We thank the Department of Science & Technology (DST), Chandigarh.

